# Thermal Adaptation of Extremozymes: Temperature-Sensitive Contact Analysis of Serine Proteases

**DOI:** 10.1101/2025.03.03.641325

**Authors:** Dulitha P. Kulathunga, Davit A. Potoyan

## Abstract

Enzyme thermal adaptation reflects a delicate interplay between sequence, structure, and dynamics of proteins, fine-tuning the catalytic activity to environmental demands. Understanding these evolutionary relationships can drive bioengineering advances, including industrial enzyme design, biocatalysts for extreme conditions, and novel therapeutics. This work explores sequence-dynamics connections in subtilisin-like serine protease homologs using a recently developed computational methodology that uses expanded ensemble simulations and temperature-sensitive contact analysis. We reveal that thermophilic enzymes achieve thermal stability through extensive salt bridges and hydrophobic networks, while psychrophilic enzymes rely on targeted interaction stability for cold adaptation. An unsupervised cluster analysis of residue conservation, flexibility, and hydrophobic interactions provides a comprehensive view of residue-specific contributions to thermal adaptation. These findings underscore the coordinated roles of conserved and variable regions in enzyme stability and offer a framework for tailoring enzymes to specific thermal properties for biotechnological applications.

**SIGNIFICANCE:** This study reveals how subtilisin-like serine proteases adapt to extreme temperatures by balancing sequence, structure, and dynamics. Through multi-ensemble simulations, we demonstrate that conserved catalytic regions and specific residue variations allow fine-tuned thermal adaptation. The analysis of simulations reveals the importance of salt bridges and hydrophobic interactions in enhancing thermophilic enzyme stability. We find that psychrophilic enzymes exhibit unique stability mechanisms for cold environments. The approach used in this study provides a comprehensive framework for understanding enzyme thermal adaptation and offers valuable insights for bioengineering enzymes with tailored properties for industrial and medical applications.

## INTRODUCTION

Extremophilic organisms thrive in environments that are incompatible with life for most species. The discovery of extremophiles in the 1960s by Brock and Freeze changed conventional ideas about the limits of life (1). Evolution has pushed extremophile genes to adapt to high or low temperatures, acidic or alkaline environments, higher salinity, and extreme pressure, making life possible from the deepest oceans to volcanic vents. Extremophiles maintain high catalytic activity under harsh thermal conditions through modified enzyme sequences, which optimize the stability activity balance for physiological temperature (2–4). This versatility enables the creation of bioreactors that operate at more efficient and energy-saving temperatures.(5)

In extremozymes, different interactions contribute uniquely to the balance of stability and flexibility found in psychrophilic and thermophilic enzymes. Salt bridges have been recognized as strong stabilizing contributors to protein structure(6, 7). Psychrophilic enzymes often have fewer salt bridges, which allows more flexibility and activity at low temperatures. In contrast, thermophilic enzymes often have more salt bridges, which increases their structural rigidity and stability against thermal denaturation(8, 9). Cationic-*π* interactions and *π*-*π* interactions also play an essential role in protein stability. Thermophilic enzymes use more aromatic interactions to increase stability at high temperatures, whereas psychrophilic enzymes favor flexibility with less aromatic interactions(8, 9). Disulfide bridges between cysteine residues further strengthen structural rigidity. Psychrophilic enzymes often contain fewer disulfide bridges, resulting in a more flexible protein structure at cold temperatures. Additional disulfide bridges in thermophilic enzymes result in a more rigid and stable structure that can survive high temperatures. However, because of the less abundant cysteine residues, the impact of disulfide bonds on thermal adaptations has been less studied (10).

Beyond these pairwise noncovalent interactions, extremozymes also adopt strategies relying on collective interactions to achieve optimal protein stability and flexibility balance. Thermophilic and psychrophilic proteins have different non-polar residue cluster properties in their protein cores. Thermophilic proteins frequently have more significant hydrophobic clusters and increased proline residues within loops(11), which improve thermal stability by encouraging tighter packing. In contrast, psychrophilic proteins may have smaller and less hydrophobic clusters,(12) but are enriched in glycines in loops, allowing for better flexibility and adaption to low temperatures(13). While comparative structural investigations have helped to shed light on differences in enzyme structures between psychrophilic and thermophilic enzymes, the evolutionary mechanisms underlying these structural modifications remain unresolved. Furthermore, more comprehensive studies that combine structural and dynamic information is lacking(14). Further research is needed to understand how sequence-specific interactions mediate thermal adaptations, especially in connection with the stability and activity of enzymes(12).

Because of the well-defined structure and availability of psychrophilic, mesophilic, and thermophilic variations, subtilisin-like serine proteases make a good model for exploring the molecular basis of thermal adaptations. In this study, we chose four homologous subtilisin enzymes that are structurally similar but tailored to four different temperature optima classified as psychrophilic (1SH7), mesophilic (1IC6), thermophilic (1THM), and extreme thermophilic enzymes (4DZT)(15–18). The cold-adapted 1SH7 was extracted from the psychrotrophic marine bacterium *Vibrio* sp. PA-44, which thrives at temperatures between 15-25°C. Mesophilic Proteinase K (1IC6), derived from *Tritirachium album limber*, exhibits optimal activity around room temperature(17). Serine Protease from *Thermoactinomyces vulgaris* (1THM), strong Cold-adapted and thermophilic proteinases demonstrated more extensive hydrogen- and ion-pair interactions than examined the protease from *Thermus aquaticus* (4DZT), a hyperthermophile with an optimum growth temperature of 70°C(18).

Subtilisin-like serine proteases have a catalytic triad consisting of serine, histidine, and aspartate residues. This triad forms the core of the active site and is responsible for its proteolytic activity(19). The catalytic mechanism of subtilisin-like serine proteases consists of two steps. The serine residue in the active site acts as a nucleophile, attacking the carbonyl carbon of the peptide bond. This results in the formation of a covalent acyl-enzyme intermediate. Then, a water molecule hydrolyzes the intermediate, releasing the cleaved peptide and restoring the free enzyme(20). This process enables effective peptide bond hydrolysis, contributing to the enzyme’s adaptability in diverse biological contexts. Arnórsdóttir et al. (2005) conducted a comparative analysis of structural features among cold-adapted (1SH7), mesophilic (1IC6), and thermophilic (1THM) subtilisin-like proteases. Despite high structural similarity, the study revealed distinct adaptations: thermophilic enzymes exhibited higher hydrophobic interactions, while cold-adapted enzymes showed greater exposure to apolar surfaces. Both cold-adapted and thermophilic proteinases demonstrated more extensive hydrogen- and ion-pair interactions compared to their mesophilic counterparts. These findings suggest that temperature adaptation in subtilisin-like proteases involves a complex interplay of structural modifications rather than a simple trade-off between flexibility and stability(17).

Sakaguchi et al. (21) further investigated the role of salt bridges in temperature adaptation by comparing the cold-adapted 1SH7 with the extreme-thermophilic 4DZT. Their study focused on introducing mutations to conserved amino acids involved in salt bridge formation in the 4DZT wild-type enzyme. They found that mutation of the highly conserved Asp183 in 4DZT led to protein structure destabilization, highlighting the significance of this particular salt bridge in maintaining the stability of 4DZT(21). These studies show the importance of specific structural features, such as hydrophobic interactions, hydrogen bonding, and salt bridges, in the temperature adaptation of subtilisin-like serine proteases. They also emphasize the complexity of enzyme adaptation to different temperature regimes, suggesting that multiple structural modifications work together to achieve optimal enzyme function across various thermal environments. Comparative investigations using molecular dynamics simulations revealed that while the overall 3D architectures are generally retained across temperature adaptations, there are considerable changes in the dynamics(22).

In this work, we illuminate the mechanistic link between sequence evolution and temperature dependence of enzyme dynamics, which has been shown to serve as a proxy for catalytic activity (23). By carrying out expanded ensemble multiple microsecond-long atomistic simulations of all four homologs at different temperatures, we capture the subtle sequence-specific dynamic differences in conformational flexibility, local and global dynamics, and stability of critical structural features such as the catalytic triad, substrate-binding regions, and salt bridges. We reveal that thermophilic enzymes achieve thermal stability through extensive salt bridges and hydrophobic networks, while psychrophilic enzymes rely on targeted interaction stability for cold adaptation. An unsupervised cluster analysis of residue conservation, flexibility, and hydrophobic interactions provides a comprehensive view of residue-specific contributions to thermal adaptation. These findings underscore the coordinated roles of conserved and variable regions in enzyme stability and offer a framework for tailoring enzymes to specific thermal properties for biotechnological applications.

## MATERIALS AND METHODS

Crystal structures of four subtilisin serine proteases (PDB IDs: 1SH7, 1IC6, 1THM, and 4DZT) were obtained from the Protein Data Bank(24). These structures were processed using PDBFixer to add missing residues and atoms. Each system was explicitly solvated in a cubic box of TIP3P water molecules with a 1 nm distance between the box edge and protein and neutralized with counter-ions to a final concentration of 100 mM NaCl. The Amber99SBILDN force field was employed for proteins and ions(25). Molecular dynamics simulations were performed using OpenMM 7.7.0.(26). After energy minimization, each enzyme was simulated independently for 4*μs* at six different temperatures ranging from 283 K to 393 K, resulting in 24 simulations of 4*μs* each. The simulations were conducted under periodic boundary conditions using the Verlet cutoff scheme. Long-range electrostatic interactions were treated using the Particle Mesh Ewald method. Temperature and pressure were maintained using the Langevin integrator and Monte Carlo barostat, respectively.

Post-simulation analyses were performed using MDAnalysis to assess structural rigidity at each temperature(27). Contact analysis was conducted using getContact and ProLIF libraries to identify critical residue network of interactions(28, 29). Temperature-sensitive contact modes were identified through Chacra analysis library(23). Graph network analysis was employed to identify temperature-stable hydrophobic networks among the enzymes, using NetworkX(30). This analysis involved constructing graphs based on hydrophobic interactions and analyzing their connectivity and stability across different temperatures. Evolutionary analysis was performed using the enzyme sequences and their respective family sequences. Multiple sequence alignment was conducted using Gremlin, and evolutionary conservation entropy was calculated using the Evol library(31, 32). Unsupervised machine learning techniques, K means clustering, were applied to compare evolutionary results with residue-wise dynamical features extracted from the simulations. These analyses were performed using scikit-learn(33). All visualizations were created using Chimera and PyMOL for structural representations(34, 35).

## RESULTS

### Sequence and Structural Comparison

Our comparative analysis of the four subtilisin-like serine proteases indicates a balance of structural conservation and divergence, showing that they adapt to various temperature environments. Despite coming from organisms with different optimal growth temperatures, these enzymes have similar overall fold, especially in key functional regions (Figure 1A). However, a closer look reveals significant structural differences, providing insights into the molecular basis of their temperature adaptations.

**Figure 1.**
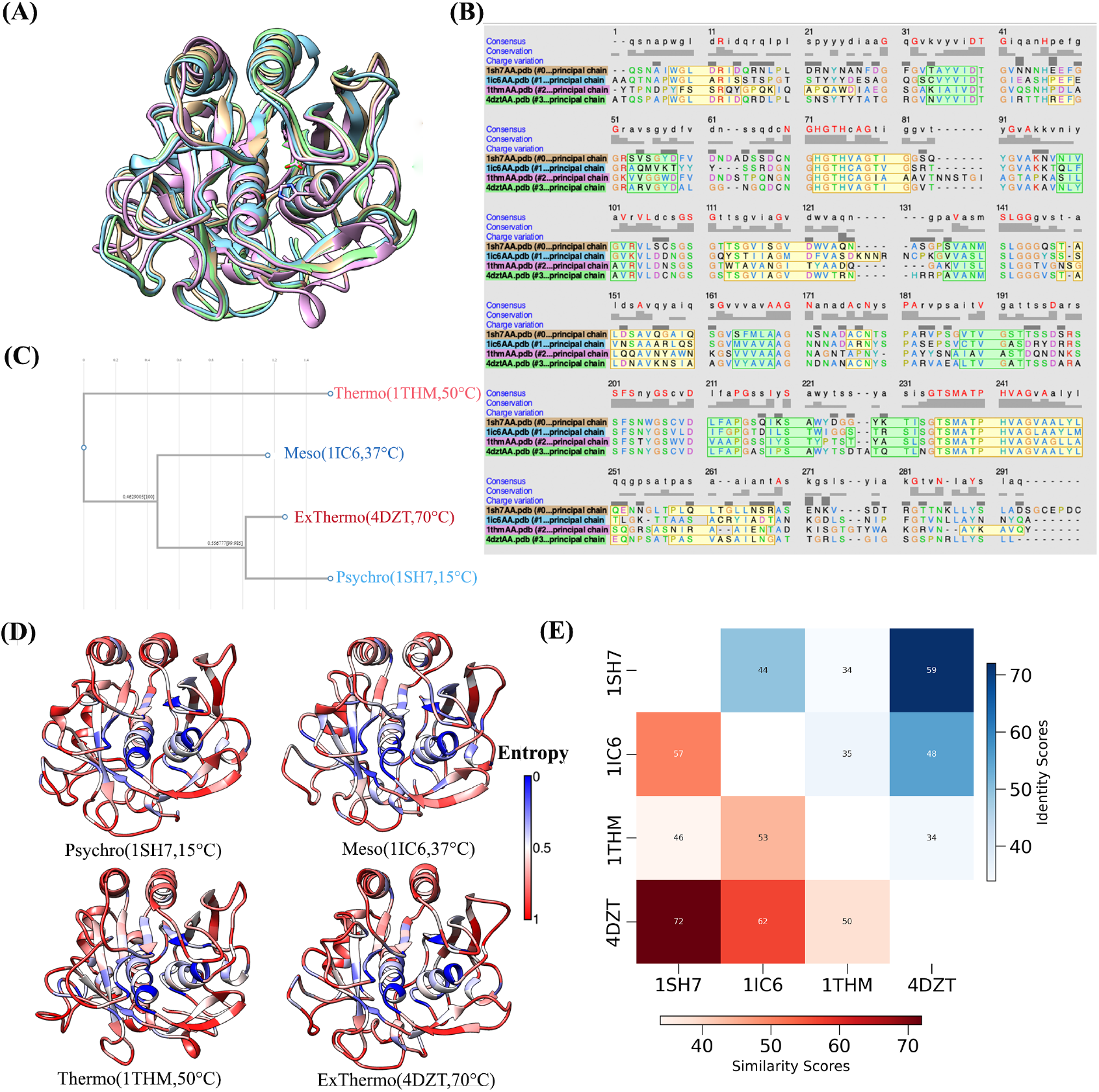
Structural and Evolutionary Analysis of Four Enzymes. A) Structural alignment of four enzymes: 1SH7 - Orange, 1IC6 - Blue, 1THM - Purple, and 4DZT - Green. B) Sequence alignment derived from the structural alignment in (A). Alpha helices and beta sheets are highlighted in green and yellow, respectively. Sequence conservation is indicated on the top of the alignment. C) Evolutionary dendrogram illustrating the phylogenetic relationships among the four enzyme sequences. D) Visualizations of the crystallographic structures of four enzymes colored by evolutionary entropy. Blue regions indicate low entropy (high conservation), while red regions show high entropy (high variability). E) Pairwise sequence identity and similarity matrix. The upper triangle displays identity scores, while the lower triangle shows similarity scores between enzyme pairs.

Figure 1A illustrates the high structural similarity among the four enzymes when aligned. The sequence alignment derived from the structural superposition (Figure 1B) provides further insights into the conservation and divergence among these enzymes. This conservation is particularly evident in the catalytic core, where most of the essential alpha helices and beta sheets are conserved throughout all four enzymes. However, some structural differences are apparent. The psychrophilic enzyme (1SH7) possesses a C-terminal extension of approximately six residues absent in the other three enzymes. This unique feature may contribute to the cold adaptation of 1SH7, possibly by providing additional flexibility at lower temperatures. Additionally, we observed slight variations in loop regions, particularly in the substrate-binding areas, which may contribute to the different thermal optima of these enzymes.

Figure1E shows the pairwise sequence similarity and identity scores, which reveal interesting patterns of proximity. The psychrophilic enzyme 1SH7 and the extreme thermophilic enzyme 4DZT have the highest sequence identity (58.7%) and similarity (72%), despite their adaption to opposite thermal extremes. This high level of conservation implies that adaptation to extreme temperatures may involve similar structural principles, with critical differences in key residues driving their distinct thermal properties. The mesophilic enzyme 1IC6 shows moderate conservation with both 1SH7 and 4DZT (identity: 44% and 47.7%; similarity: 57.4% and 61.8%, respectively), positioning it as an intermediate variant.

The thermophilic enzyme 1THM consistently exhibits the lowest identity and similarity scores across all sequences (ranging from 33.9% to 35% identity and 46.4% to 53.2% similarity), indicating it is the most divergent among the four. The distinctive trait of 1THM suggests it may have evolved unique strategies for thermostability, possibly representing an alternative evolutionary path within the subtilisin family. These observations suggest the complexity of protein thermal adaptation, highlighting that while overall sequence conservation may be high, subtle differences in key regions significantly affect an enzyme’s thermal properties.

The evolutionary dendrogram (Figure 1C) derived from the sequences of these four subtilisin-like serine proteases provides further insights that corroborate and extend our understanding of their evolutionary relationship. The tree topology closely matches the patterns observed in the pairwise sequence comparisons, visually representing the complex evolutionary history of these thermally diverse enzymes. The dendrogram shows the psychrophilic enzyme (1SH7) and the extreme thermophilic enzyme (4DZT) clustering closely together, forming a distinct clade. This close evolutionary relationship mirrors their high sequence identity and similarity scores (58.7% and 72%, respectively). This clustering suggests that these enzymes likely share a recent common ancestor, with thermal specialization occurring through a relatively small number of critical mutations rather than whole sequence divergence. This challenges simplistic models of thermal adaptation and hints at the possibility of shared molecular mechanisms underlying cold and heat adaptation in this enzyme family.

The mesophilic enzyme (1IC6) appears as a separate branch in the dendrogram, positioned between the 1SH7-4DZT clade and the thermophilic enzyme (1THM). This intermediate placement is consistent with its moderate sequence similarity to both extremophilic enzymes and aligns with its adaptation to intermediate temperatures. This suggests that 1IC6 may retain ancestral features that allow for more straightforward adaptation to both temperature extremes, potentially representing an evolutionary midpoint. The position of the thermophilic enzyme (1THM), which emerges as the most divergent, occupies a separate and distant branch. This distinct evolutionary position matches the consistently low sequence identity and similarity scores observed for 1THM. The substantial evolutionary distance between 1THM and the other enzymes, including the extreme thermophilic 4DZT, suggests that 1THM may have evolved unique strategies for thermostability. This divergence raises intriguing questions about the multiple potential evolutionary pathways to thermal stability within the subtilisin family.

To better understand the evolutionary conservation patterns of these subtilisin-like serine proteases, we calculated informational entropy as a residue conservation score using multiple sequence alignments (MSAs) from their respective protein families. Each enzyme’s MSA contained around 3000 homologous sequences, presenting a solid statistical foundation for the investigation. The Shannon entropy was calculated using the Evol library, with low entropy indicating highly conserved residues among the sequence family and high entropy signifying highly variable residues. The normalized entropy scores were then projected onto the respective protein structures (Figure 1D), revealing notable conservation across the enzymes. Notably, the core residues of all four enzymes, particularly those encompassing the catalytic triad and substrate-binding pocket, exhibited remarkably low entropy scores, indicating high evolutionary conservation. This conservation emphasizes the critical functional importance of these core residues in maintaining the enzymes’ catalytic activity across diverse temperature environments.

In contrast, surface-exposed residues and terminal regions displayed higher entropy scores, suggesting greater evolutionary flexibility in these areas. As evident in Figure 1D, most internal residues are conserved, while most mutations occur in all four enzymes’ surface and terminal residues. This trend is consistent with the idea that functionally important core regions are more conserved. Still, surface residues are more responsive to alterations that contribute to specific adaptations, such as thermal stability or flexibility. Moreover, this entropy-based analysis provides a quantitative measure of evolutionary conservation, which improves our understanding of the individual residues and regions subject to selective pressure during the thermal adaptation of these enzymes.

### Structural Flexibility

To investigate the dynamic behavior of these subtilisin-like serine proteases across different thermal environments, we analyzed the root mean square fluctuations (RMSF) of C*α* atoms derived from molecular dynamics simulations at six different temperatures for each enzyme (Figure 2 A). This analysis provides crucial insights into these enzymes’ structural flexibility and rigidity, pivotal in the activity-stability trade-off observed in extremozymes.

As expected, the RMSF profiles reveal distinct patterns of flexibility among the psychrophilic (1SH7), mesophilic (1IC6), and thermophilic (4DZT, 1THM) enzymes, reflecting their adaptations to different thermal environments. The extreme thermophilic enzyme 4DZT exhibits remarkably low fluctuations across its backbone as temperatures increase, indicative of its enhanced structural rigidity. This rigidity is likely a key factor in maintaining stability at elevated temperatures. However, 4DZT does show increased flexibility in specific regions, notably residues 100-104 and 130-140, which are proximal to the substrate-binding site. This localized flexibility at higher temperatures may be crucial for maintaining catalytic activity under extreme conditions.

In contrast, the psychrophilic enzyme 1SH7 demonstrates higher overall fluctuations in response to temperature changes, particularly in its terminal regions. This increased flexibility is consistent with the general observation that cold-adapted enzymes require enhanced conformational plasticity to achieve catalytic efficiency at low temperatures. Interestingly, 1SH7 maintains some core rigidity despite its cold adaptation compared to the mesophilic enzyme 1IC6, especially at higher temperatures. This unexpected feature may be related to the close evolutionary relationship between 1SH7 and 4DZT observed in our sequence analysis, suggesting a complex interplay between evolutionary history and functional adaptation.

The mesophilic enzyme 1IC6 shows high fluctuations in core regions at very high temperatures, surpassing 1SH7 and 4DZT in these areas. This increased flexibility in core regions at extreme temperatures may explain the limited thermal stability of mesophilic enzymes compared to their thermophilic counterparts. Notably, the thermophilic enzyme 1THM shows increased flexibility in specific regions, including the N-terminal area, residues 55-70 near the catalytic residue His70, and the substrate-binding region (residues 104-108). This localized flexibility and overall structural rigidity may represent a fine-tuned balance that allows these enzymes to maintain stability and activity at high temperatures.

To further understand the temperature-dependent flexibility of these subtilisin-like serine proteases, we calculated the standard deviation of RMSF across different temperatures for each residue, which served as a temperature sensitivity score. This analysis provides an advanced view of how various regions of each enzyme respond to temperature changes. Figure 2B shows a heatmap that compares all four enzymes based on their structural sequence alignment, providing a comprehensive view of their temperature-sensitive regions.

**Figure 2.**
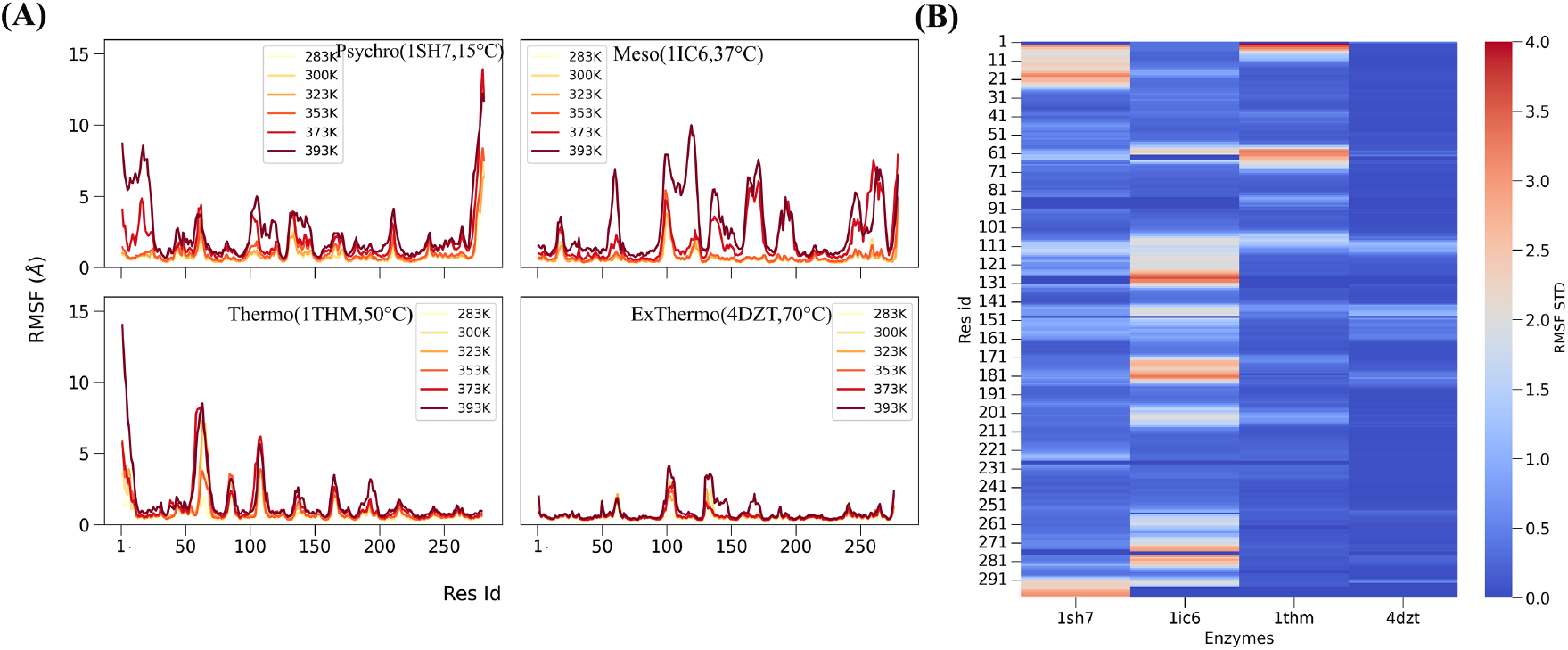
Temperature-Dependent Structural Flexibility Analysis. A) RMSF plots for each enzyme across six temperatures. Subplots are arranged: top left - 1SH7, top right - 1IC6, bottom left - 1THM, bottom right - 4DZT. Each line represents RMSF at a different temperature, illustrating how each enzyme’s structural flexibility changes with temperature. B) Residue-wise comparison of temperature sensitivity scores for the four enzymes. Scores are calculated from the standard deviation of RMSF values across six temperatures for each residue in each enzyme. Higher scores indicate greater temperature-induced flexibility.

Figure 2B reveals distinct flexibility and temperature sensitivity patterns among the four enzymes, extending our previous RMSF observations. The extreme thermophilic enzyme 4DZT exhibits the lowest overall RMSF variance, indicating its remarkable stability across a wide temperature range. This stability is likely a key factor in its ability to maintain catalytic function under extreme thermal conditions. In contrast, the mesophilic enzyme 1IC6 displays localized regions of high RMSF variance, indicating areas of the protein that are particularly sensitive to temperature changes. These flexible regions may assist the enzyme in adapting to moderate shifts in temperature but also explain its limited performance at extreme temperatures.

The psychrophilic enzyme 1SH7 has a distinct pattern of flexibility, with substantial RMSF variance localized at the N- and C-termini. This terminal flexibility may be crucial to sustaining catalytic activity at low temperatures by allowing for essential conformational changes. Localizing flexibility to the terminal regions while keeping the core relatively stable could be an evolutionary strategy for balancing low-temperature activity with structural integrity. The thermophilic enzyme 1THM, while generally showing lower RMSF variance than 1IC6 and 1SH7, exhibits some regions of increased temperature sensitivity compared to 4DZT. This intermediate pattern of flexibility aligns with its moderate thermostability and suggests a balanced approach to maintaining stability and necessary flexibility for catalytic function at elevated temperatures.

These findings highlight the complex relationship between local flexibility, global stability, and enzyme temperature adaptation. The extreme thermophilic 4DZT appears to have evolved a structure that is uniformly resistant to temperature-induced fluctuations, while the psychrophilic 1SH7 and mesophilic 1IC6 show more localized areas of flexibility that may be crucial for their function in their respective thermal environments. The thermophilic 1THM represents the necessity for overall stability while retaining flexibility in critical regions. Our findings provide solid support for the activity-stability tradeoff in extremozymes.

### Non-covalent interactions

Non-covalent interactions are essential for maintaining enzymes’ structural stability and functional attributes across diverse thermal conditions. Our analysis of contact frequencies at different temperatures, specifically examining salt bridges, hydrophobic interactions, and other non-covalent interactions, revealed distinct patterns that provided insight into the molecular-level mechanisms of thermal adaptation in these subtilisin-like serine proteases.

The contact frequency heat maps (SI Figure 1) demonstrate differences in salt bridge stability among the four enzymes. The extreme thermophilic enzyme 4DZT has many stable salt bridges at all temperatures screened, also shared by the thermophilic enzyme 1THM. Stable salt bridges are expected to contribute significantly to their structural rigidity and thermal stability at high temperatures. Although 1THM and 4DZT enzymes have many stable salt bridges contributing to their thermal stability, the specific salt bridge interactions vary considerably. Thermophilic 1THM has a unique distribution of salt bridges that, while contributing to its overall stability, may reflect a divergence in their evolutionary paths despite their shared thermophilic nature. In contrast, the mesophilic enzyme 1IC6 displays the most unstable salt bridge interactions, suggesting a more dynamic structure that may facilitate activity at moderate temperatures but compromises stability at higher temperatures.

Despite its ability to adapt to lower temperatures, the cold-adapted enzyme 1SH7 has more stable salt bridges than the mesophilic 1IC6. This finding may be linked to the close evolutionary relationship between 1SH7 and 4DZT observed in our earlier sequence analysis. It suggests that cold adaptation in 1SH7 may involve a more complex strategy than simply reducing overall structural rigidity. We identified several highly conserved salt bridges across all four enzymes, such as ARG10 – ASP183 in 1SH7 (and its equivalents in other enzymes). The conservation of these interactions across enzymes adapted to different thermal environments suggests their fundamental importance in maintaining the overall protein structure. Additionally, we observed salt bridges common to 1SH7 and 4DZT, such as ARG95 – ASP56, which may represent ancestral features retained aside from divergent thermal adaptations.

Our analysis (SI Figure 2,3) did not reveal any significant cationic-*π* or *π*-*π* interactions among these four enzymes that could be attributed to their temperature adaptations. This suggests that these types of interactions may play a less significant role in determining the thermal properties of these particular enzymes.

### Hydrophobic Networks

We conducted an in-depth analysis of hydrophobic interactions using graph theory approaches. We calculated average hydrophobic contact frequencies across temperatures and constructed graph networks for each enzyme, focusing on stable contacts with frequencies higher than 0.75. In these networks, hydrophobic interactions are represented as edges, with residues as nodes and edge width indicating the average contact frequency. Among the many measures calculated, closeness centrality (CC) was revealed as a particularly informative metric to evaluate the temperature sensitivity of hydrophobic networks. For a residue *i*, closeness centrality (*C*_*i*_) was computed as the inverse of the sum of shortest path lengths to all other residues *j*, normalized by the total number of residues (*n* – 1):

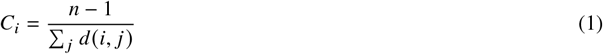

where *d(i, j)* is the shortest path length between residues *i* and *j* (30). CC analysis revealed key residues for maintaining structural stability across various thermal conditions and how the dynamic interplay of hydrophobic interactions contributes to thermal stability and functional flexibility throughout the temperature spectrum.

Figure 3 illustrates the distribution of CC scores across the four enzymes, revealing a clear trend from psychrophilic to extreme thermophilic adaptations. To visualize these key interactions, we extracted the top 20% CC residue interactions as a sub-network and projected them onto the PDB structures (Figure 3A). The extreme thermophilic enzyme 4DZT and the thermophilic 1THM exhibit significantly higher residues with high CC values than the psychrophilic 1SH7. Notably, 4DZT shows the highest overall CC values, indicating a well-connected hydrophobic core. This suggests that the extreme thermophilic enzyme has evolved a more integrated and stable hydrophobic network to maintain structural integrity at high temperatures. Numerous high CC residues distributed throughout the protein structure in 4DZT and 1THM suggest that thermophilic enzymes rely on a well-integrated hydrophobic core to maintain their structural rigidity under extreme conditions.

**Figure 3.**
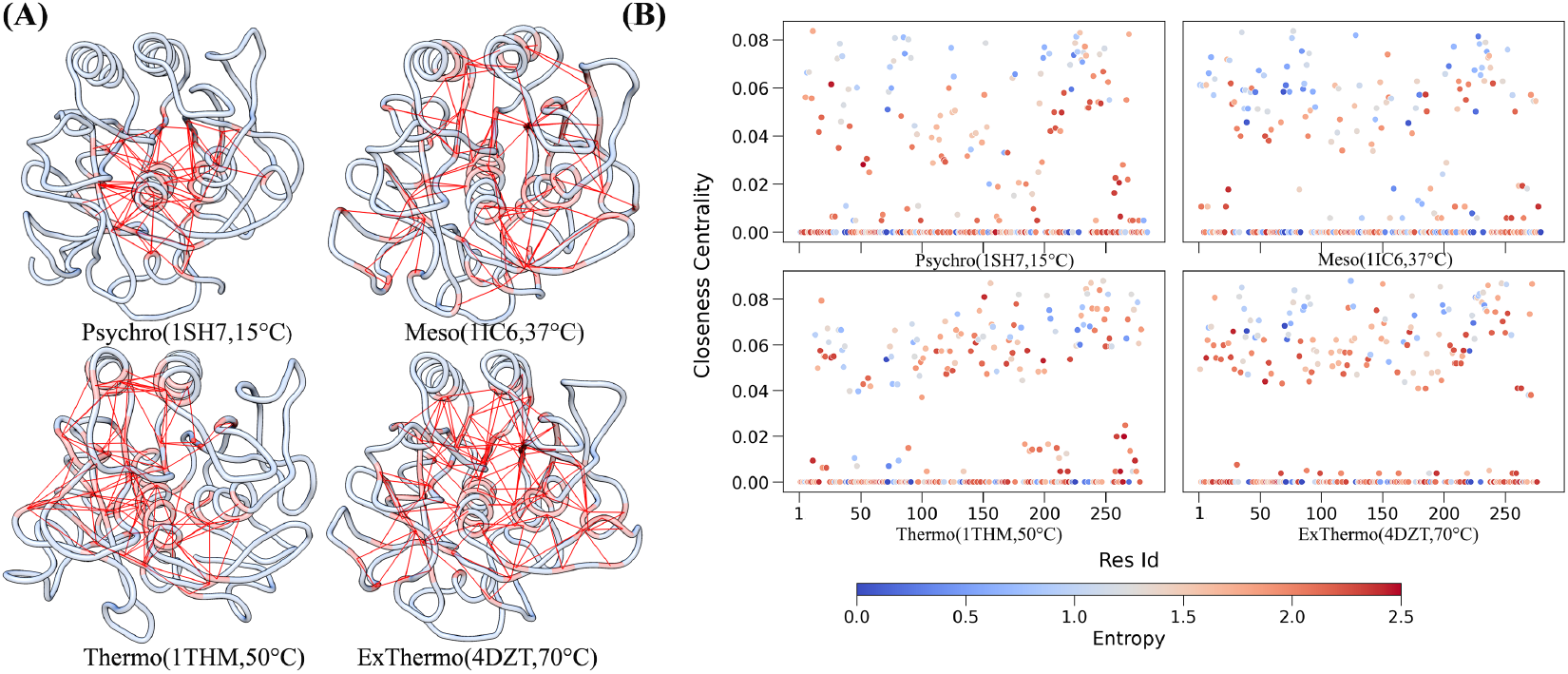
Analysis of Hydrophobic Networks. A) PDB structure visualizations highlight the top 20% residues with the highest closeness centrality and their hydrophobic networks for each enzyme. Structures are arranged as follows: top left - 1SH7, top right - 1IC6, bottom left - 1THM, bottom right - 4DZT. These visualizations demonstrate the spatial distribution of key hydrophobic networks within each enzyme’s structure. B) Scatter plots of closeness centrality scores for hydrophobic networks as a function of residue ID for each enzyme. Points are colored according to evolutionary residue entropy, with blue indicating low entropy (high conservation) and red indicating high entropy (high variability). Plots are arranged in the same order as in panel A: top left - 1SH7, top right - 1IC6, bottom left - 1THM, bottom right - 4DZT.

Interestingly, the psychrophilic enzyme 1SH7 displays higher CC values for some residues, showing that 1SH7 has a more complex adaptation strategy than initially anticipated. This finding suggests that cold adaptation in 1SH7 involves a delicate balance between maintaining structural stability and allowing for the increased flexibility required for catalytic activity at low temperatures. Some high CC residues in 1SH7, even surpassing those in 1IC6, indicate that psychrophilic adaptation is not simply a matter of reduced hydrophobic interactions but a strategic redistribution of these interactions. The mesophilic enzyme 1IC6, with its intermediate hydrophobic network characteristics, represents a more flexible structure capable of functioning across a moderate temperature range. Its less extensive high CC network than thermophilic enzymes likely contributes to its limited stability at extreme temperatures. Still, it may allow for greater conformational flexibility necessary for its biological role. Overlaying the evolutionary entropy data on the CC scatter plot (Figure 3B) reveals that most high closeness centrality residues are conserved across all four enzymes. This conservation explains the fundamental importance of these hydrophobic interactions in maintaining the core structure and fold of the enzymes, regardless of their thermal adaptation. The conserved core likely serves as a foundation upon which specific thermal adaptations are built by modulating peripheral hydrophobic interactions. Rather than simple increases or decreases in overall hydrophobicity, our analysis reveals that adaptation involves strategic changes in the distribution and connectivity of hydrophobic residues.

### Chemically Accurate Contact Response Analysis

To better understand the non-covalent interactions of thermal adaptation in subtilisin-like serine proteases, we used Chemically Accurate Contact Response Analysis (ChACRA) to identify important contact pairs required for temperature responsiveness (23). This method uses contact frequencies generated through the getContact library and applies principal component analysis (PCA) to identify unique temperature-sensitive contact modes. Our analysis focused on the first two principal components (PCs), which account for the most significant variations in contact response across temperatures. PC1 mostly portrays melting contacts, which weaken or break as temperatures rise. In contrast, PC2 relates to the formation of contacts, which indicate interactions that strengthen or arise as temperatures increase (23).

Figure 4 illustrates the distribution of contact modes across the temperature spectrum, from psychrophilic to extreme thermophilic enzymes. We observed a clear separation of contact modes as we progressed from cold-adapted to heat-adapted enzymes. In the psychrophilic enzyme (1SH7) and the mesophilic enzyme (1IC6), the majority of contacts are captured by the melting mode (PC1). This suggests that these enzymes, particularly the psychrophilic variant, exhibit highly dynamic behavior with contacts continuously forming and breaking, consistent with the flexibility required for catalysis at lower temperatures. In contrast, the thermophilic enzymes display a more rigid behavior. While some regions show contact melting, others demonstrate the formation of new, essential contacts. This behavior suggests a sophisticated mechanism for maintaining structural integrity at elevated temperatures. The formation of new contacts in specific regions likely contributes to the enhanced rigidity characteristic of thermophilic enzymes, which is crucial for their stability and function at high temperatures.

### Temperature-sensitive residue clustering

To gain a holistic understanding of the residue-level features contributing to thermal adaptation, we employed unsupervised clustering techniques to identify essential residue clusters based on evolutionary conservation, flexibility, and hydrophobic interactions. This approach allows us to integrate multiple aspects of protein structure and dynamics, providing a more comprehensive view of the residue-level thermal adaptation in subtilisin-like serine proteases. We used evolutionary entropy, RMSF standard deviation, and closeness centrality scores as features for clustering each enzyme’s residues. After determining the optimal number of clusters using the elbow method for the within-cluster sum of squares, we applied k-means clustering with four clusters. Clustering results are shown in Figure 5 as a bivariate comparison of RMSF and closeness centrality against evolutionary entropy.

**Figure 4.**
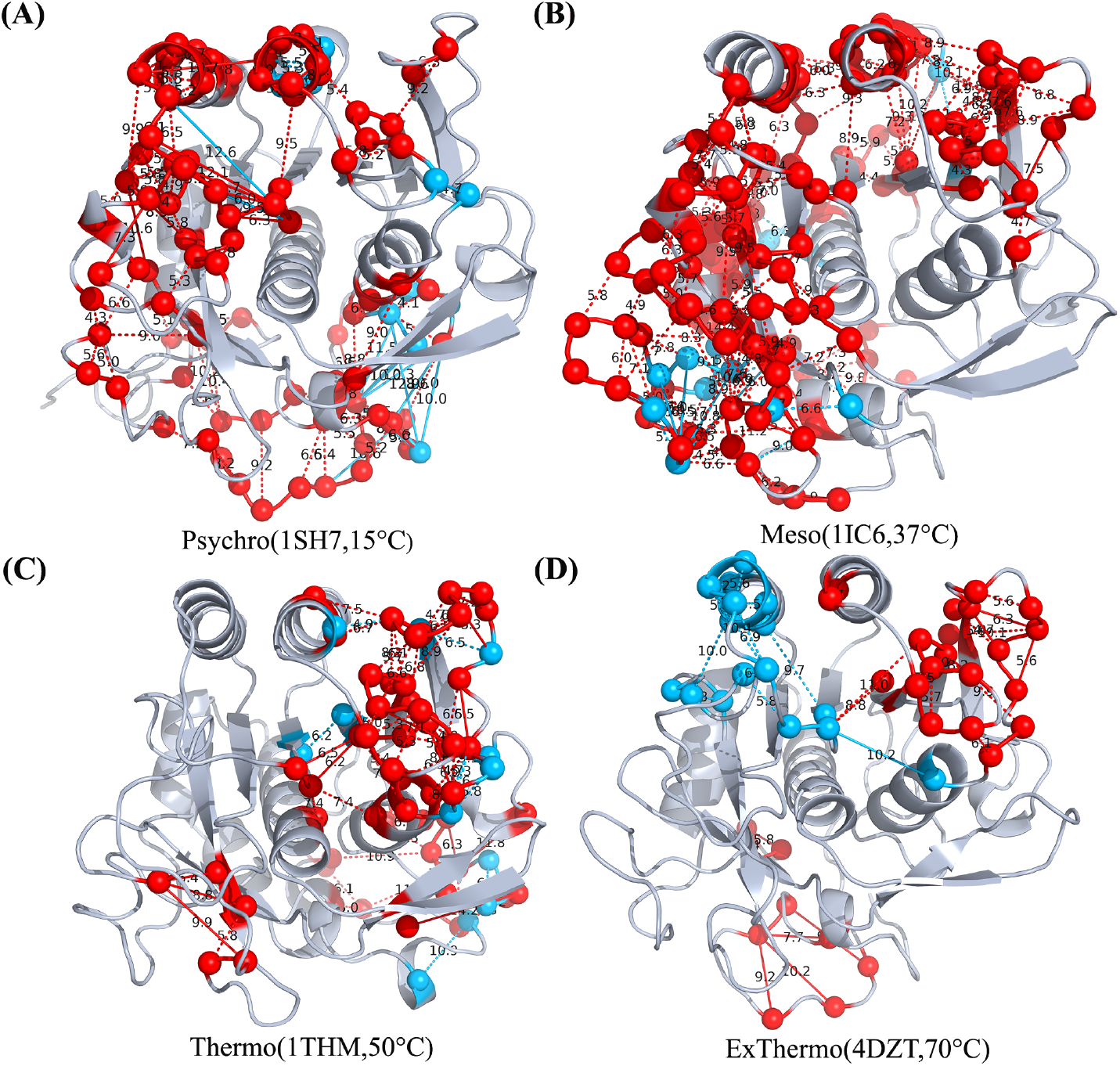
Chemically Accurate Contact Response Analysis reveals thermal modes containing the contacts that are most sensitive to temperature. A-D) Top residue contacts corresponding to the first two thermal modes (PC1 in red and PC2 in blue) for (A) 1SH7, (B) 1IC6, (C) 1THM, and (D) 4DZT protein structures. Lines indicate significant interactions between residue pairs. As thermal stability increases from psychrophilic to extreme thermophilic proteins, a clear separation of melting (primarily captured in psychrophilic structures) and forming interactions becomes evident. This separation reflects the distinct dynamical behaviors and flexibility characteristics across the thermal adaptation spectrum.

**Figure 5.**
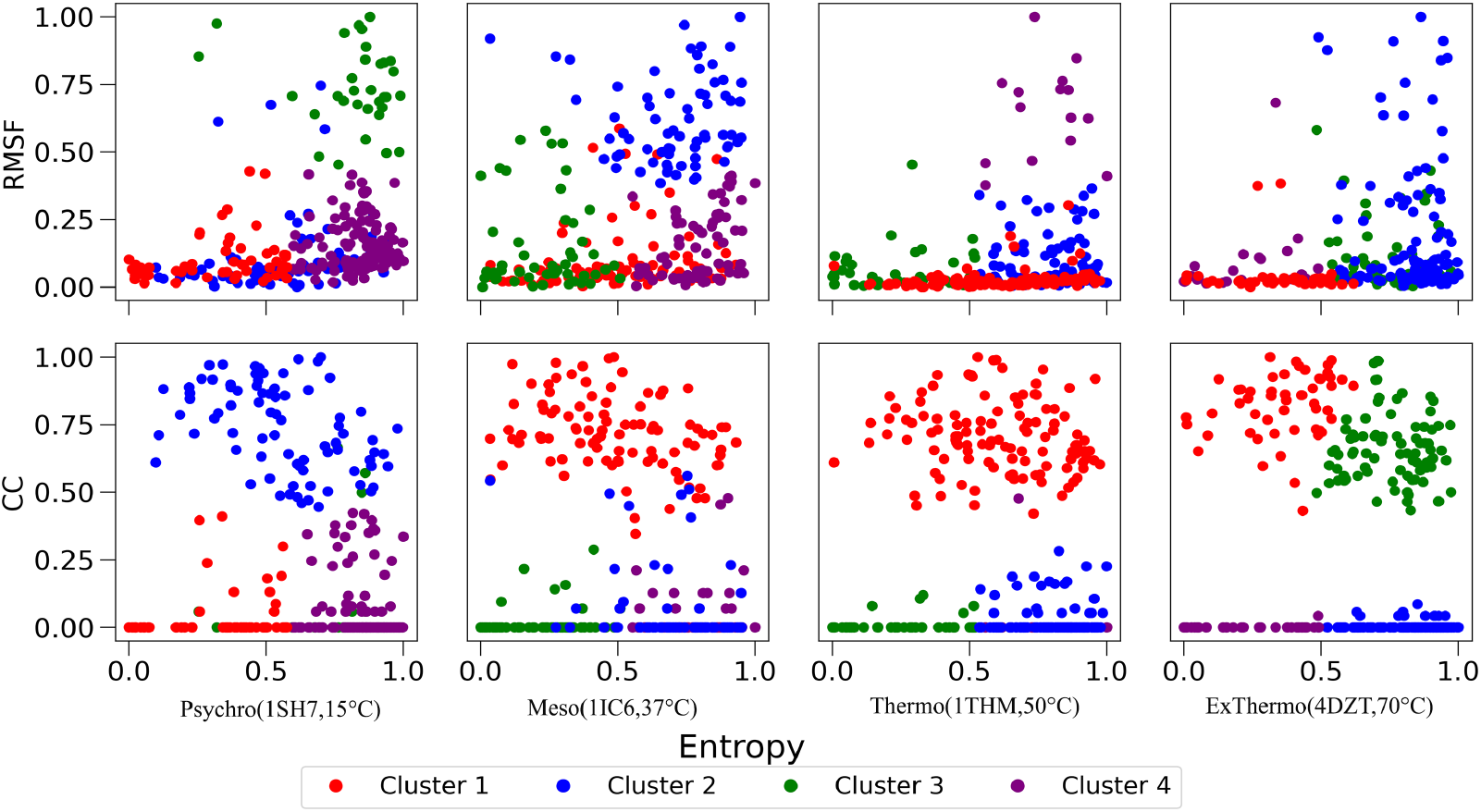
Residue cluster projections in two-dimensional comparisons of Root Mean Square Fluctuation (RMSF) and Closeness Centrality (CC) against Evolutionary Entropy for four subtilisin-like serine proteases. Each column represents one enzyme: (A) 1SH7, (B) 1IC6, (C) 1THM, and (D) 4DZT. The first row shows RMSF vs. Evolutionary Entropy, and the second row shows CC vs. Evolutionary Entropy. Residues are grouped into four distinct clusters, represented by different colors within each enzyme. These projections reveal the interplay between structural dynamics, network centrality, and evolutionary conservation across different residues of each enzyme.

In the psychrophilic enzyme 1SH7, we identified four distinct clusters with unique characteristics. Cluster 3 is characterized by low conservation and high flexibility, which is likely crucial in the enzyme’s cold adaptation. This cluster’s low closeness centrality values suggest that these residues are less involved in hydrophobic interactions, potentially contributing to the flexibility required for low-temperature activity. Conversely, Cluster 2 comprises residues with high closeness centrality, indicating important hydrophobic interactions. These residues show lower flexibility compared to Cluster 3, suggesting a role in maintaining some structural stability while allowing for overall enzyme flexibility. Clusters 1 and 4 consist of less flexible and less hydrophobic residues, with Cluster 1 showing higher conservation, indicating a more stable core necessary for maintaining basic structural integrity.

The mesophilic enzyme 1IC6 exhibits a different clustering pattern. Cluster 2 contains less conserved and highly flexible residues, similar to Cluster 3 in 1SH7. In contrast, Cluster 1 represents high closeness centrality residues with mixed conservation, potentially indicating adaptable hydrophobic interactions. Clusters 3 and 4 show lower closeness centrality and flexibility, differing primarily in their evolutionary conservation.

In the thermophilic enzymes, distinct clustering patterns reflect their adaptation to high temperatures. For 1THM, Cluster 4 contains a small number of flexible, less conserved residues, while Cluster 1 consists of high closeness centrality residues crucial for thermostability. Clusters 2 and 3 are characterized by less flexibility, with Cluster 3 showing higher conservation. The extreme thermophilic enzyme 4DZT displays different residue clusters than the other enzymes. Cluster 2 contains the most flexible residues with high evolutionary entropy, while Clusters 1 and 3 have high closeness centrality, with Cluster 1 exhibiting the highest conservation. Cluster 4 shows less flexibility, lower entropy, and lower closeness centrality.

This unsupervised clustering analysis provides several key insights into the molecular basis of thermal adaptation. First, identifying distinct, flexible clusters in each enzyme, particularly prominent in the psychrophilic 1SH7, explains the importance of localized flexibility in cold adaptation. The gradual reduction of this cluster size in mesophilic and thermophilic enzymes illustrates a clear trend in the trade-off between flexibility and stability across thermal adaptations. Second, the varying distribution of high closeness centrality clusters across the enzymes reveals how hydrophobic interactions are optimized for different thermal environments. The extensive high CC clusters in thermophilic enzymes, particularly 4DZT, highlight the crucial role of hydrophobic interactions in maintaining stability at high temperatures.

The integration of evolutionary entropy in our analysis reveals that thermal adaptation involves a complex interplay between conserved and variable regions. Highly conserved clusters in all enzymes likely represent core functional or structural elements, while variable clusters may be key to specific thermal adaptations. The clustering results visually demonstrate how each enzyme achieves a unique balance between flexible and rigid regions tailored to its thermal niche. This balance is particularly evident in the mesophilic 1IC6, which shows a more even distribution of flexibility across clusters. By identifying specific residue clusters associated with different aspects of thermal adaptation, this analysis provides valuable targets for protein engineering efforts. Modifying residues within these clusters could predictably alter an enzyme’s thermal properties.

## CONCLUSION

This work reveals how subtilisin-like serine proteases’ sequence, structure, and dynamics synergize to enable thermal adaptation across various environments. Structural comparison shows that the core of catalytic regions for all protease homologs is well conserved. Despite their distinct thermal properties, the psychrophilic enzyme 1SH7 and the extreme thermophilic enzyme 4DZT exhibit higher sequence similarity. This finding suggests that similar evolutionary strategies have been used for Serine protease adaptation to extreme cold and hot temperatures. Additionally, the evolutionary divergence of thermophilic enzyme 1THM points to multiple evolutionary pathways to achieve thermal stability within this enzyme family.

We identified notable local differences in conformational dynamics, further illuminating differences between enzymes and showing that thermophiles have greater core rigidity while displaying localized flexibility near catalytic sites. Thus, a subtle balance between stability and conformational flexibility is required for catalysis. Increased flexibility in mesophilic enzymes’ core and psychrophilic enzymes’ terminal regions at extreme temperatures may explain their reduced thermal stability. Furthermore, the unusual stability of 1SH7 homolog relative to 1IC6 highlights a tradeoff between activity and stability in extreme cold, offering further insights into the close evolutionary connection between 1SH7 and 4DZT.

Our exploration of non-covalent interactions, particularly salt bridges and hydrophobic networks, offered additional insights into these enzymes’ structural integrity strategies. Thermophilic enzymes demonstrated more robust and extensive networks of these interactions, contributing to their enhanced thermal stability. Interestingly, the psychrophilic enzyme revealed unexpected stability in specific interactions, hinting at a nuanced approach to cold adaptation that balances flexibility with structural integrity. Unsupervised clustering analysis, which integrated conservation, flexibility, and hydrophobic interactions, provided a holistic perspective on the residue groups contributing to thermal adaptation. This analysis uncovered distinct clusters linked to various thermal properties, demonstrating that thermal adaptation arises from a coordinated effort involving the enzyme structure’s conserved and variable regions. Overall, our findings indicate that enzyme temperature adaptation is a multifaceted process that requires the fine-tuning of various structural and dynamic features. The insights gained from this study deepen our understanding of how subtilisin-like serine proteases adapt to diverse thermal environments while also providing a methodological framework for exploring thermal adaptation in other enzyme families.

## AUTHOR CONTRIBUTIONS

DPK and DAP designed the research. DPK carried out all simulations and analyses. DPK and DAP wrote the manuscript

## ACKNOWLEDGMENTS

This work was supported by funds from the National Institute of General Medical Sciences with grant no. R35 GM138243 awarded to DAP.

## SUPPLEMENTARY MATERIAL

An online supplement to this article can be found by visiting BJ Online at http://www.biophysj.org.

