## Supporting Information for "Thermal Adaptation of Extremozymes: Temperature-Sensitive Contact Analysis of Serine Proteases"

### 1 SUPPLEMENTARY FIGURES.

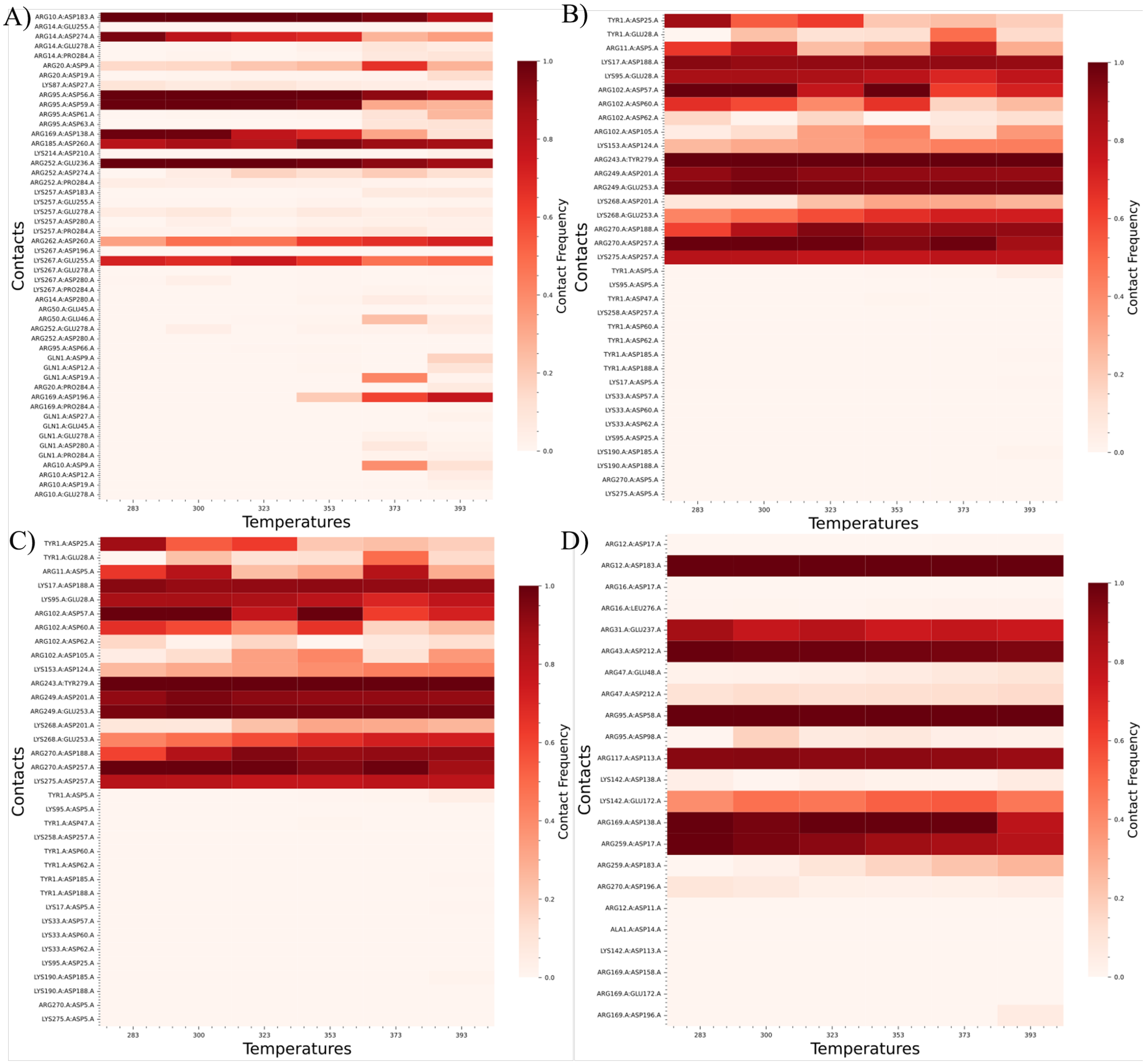


Figure 1: **Contact frequency heatmaps for cationic interactions across six temperatures** (283K to 393K). (A) Top left: Contact frequency heatmap for psychrophilic 1SH7. (B) Top right: Contact frequency heatmap for mesophilic 1IC6. (C) Bottom left: Contact frequency heatmap for thermophilic 1THM. (D) Bottom right: Contact frequency heatmap for extreme thermophilic 4DZT. Each heatmap displays the contact frequency of cationic interactions at six different temperatures: 283K, 300K, 323K, 353K, 373K, and 393K. The color scale ranges from light to dark red, with darker red indicating higher contact frequency. The most stable contacts, represented by dark red across all temperatures, demonstrate high contact frequency maintained throughout the temperature range.

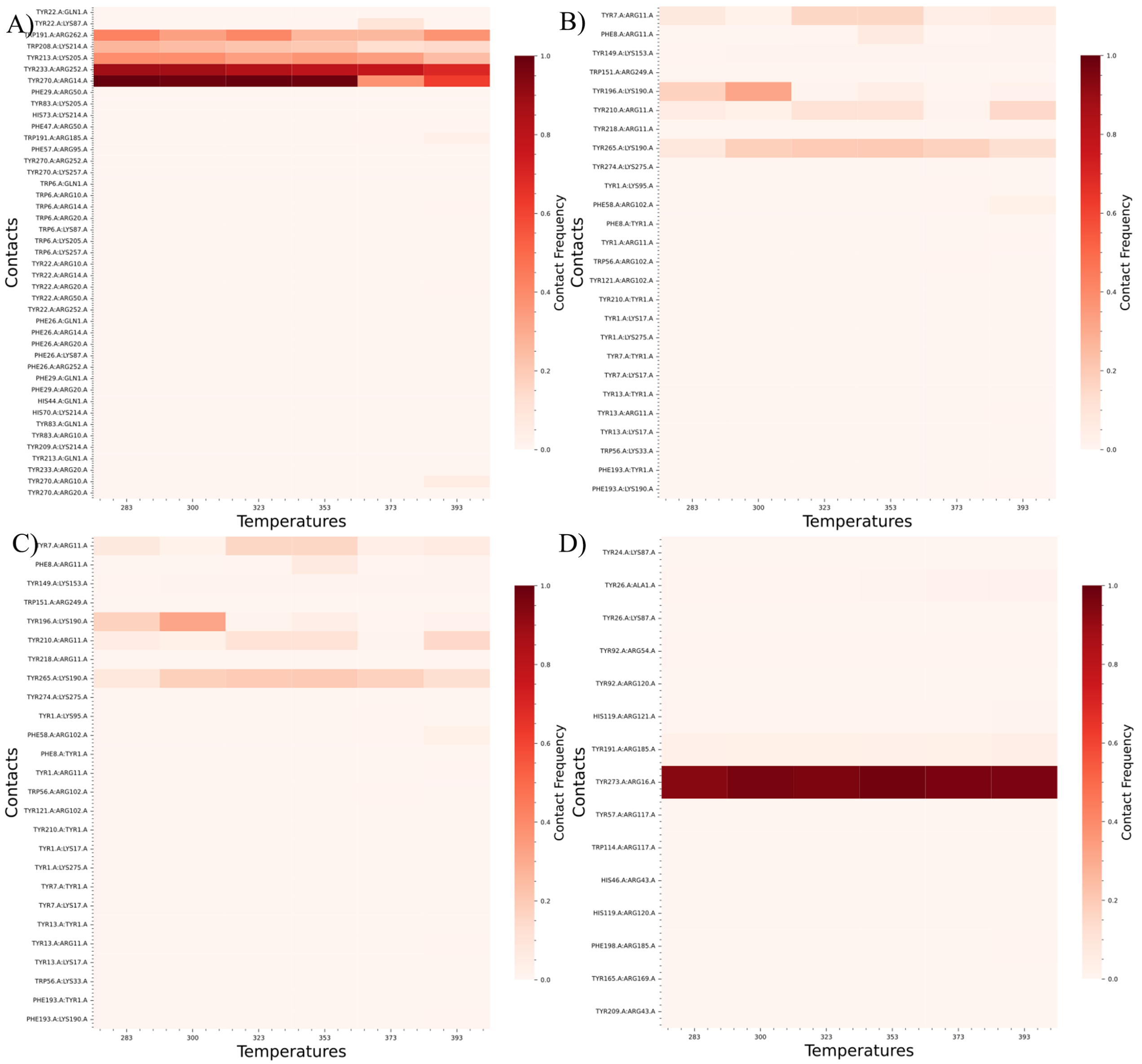


Figure 2: **Contact frequency heatmaps for pi-cationic interactions across six temperatures** (283K to 393K). (A) Top left: Contact frequency heatmap for psychrophilic 1SH7. (B) Top right: Contact frequency heatmap for mesophilic 1IC6. (C) Bottom left: Contact frequency heatmap for thermophilic 1THM. (D) Bottom right: Contact frequency heatmap for extreme thermophilic 4DZT. Each heatmap displays the contact frequency of cationic interactions at six different temperatures: 283K, 300K, 323K, 353K, 373K, and 393K. The color scale ranges from light to dark red, with darker red indicating higher contact frequency. The most stable contacts, represented by dark red across all temperatures, demonstrate high contact frequency maintained throughout the temperature range.

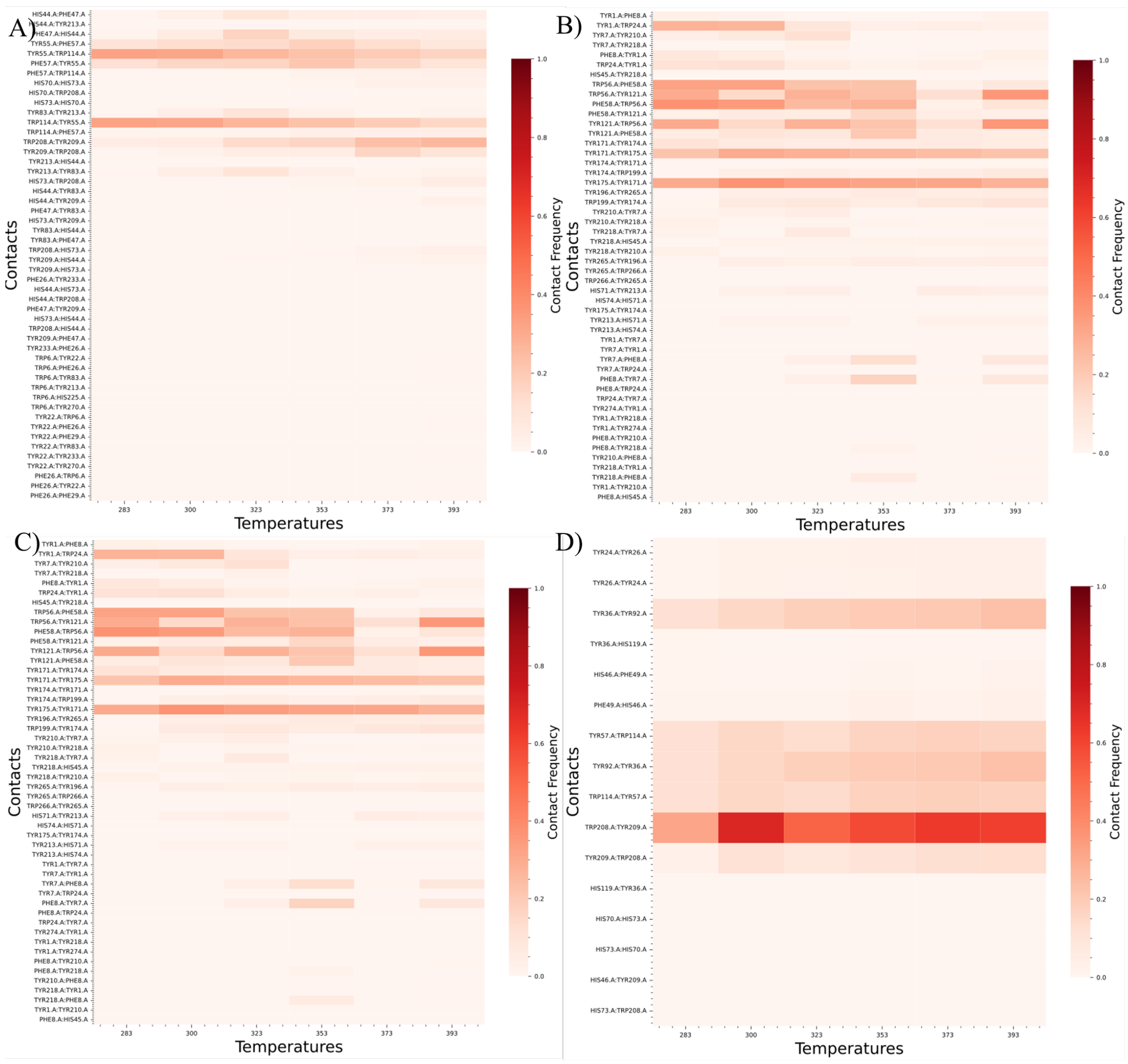


Figure 3: **Contact frequency heatmaps for pi-stacking interactions across six temperatures** (283K to 393K). (A) Top left: Contact frequency heatmap for psychrophilic 1SH7. (B) Top right: Contact frequency heatmap for mesophilic 1IC6. (C) Bottom left: Contact frequency heatmap for thermophilic 1THM. (D) Bottom right: Contact frequency heatmap for extreme thermophilic 4DZT. Each heatmap displays the contact frequency of cationic interactions at six different temperatures: 283K, 300K, 323K, 353K, 373K, and 393K. The color scale ranges from light to dark red, with darker red indicating higher contact frequency. The most stable contacts, represented by dark red across all temperatures, demonstrate high contact frequency maintained throughout the temperature range.
